# Larger and denser: an optimal design for surface grids of EMG electrodes to identify greater and more representative samples of motor units

**DOI:** 10.1101/2023.02.18.529050

**Authors:** Arnault H. Caillet, Simon Avrillon, Aritra Kundu, Tianyi Yu, Andrew T.M. Phillips, Luca Modenese, Dario Farina

## Abstract

The spinal motor neurons are the only neural cells whose individual activity can be non-invasively identified. This is usually done using grids of surface electromyographic (EMG) electrodes and source separation algorithms; an approach called EMG decomposition. In this study, we combined computational and experimental analyses to assess how the design parameters of grids of electrodes influence the number and the properties of the identified motor units. We first computed the percentage of motor units that could be theoretically discriminated within a pool of 200 simulated motor units when decomposing EMG signals recorded with grids of various sizes and interelectrode distances (IED). Increasing the density, the number of electrodes, and the size of the grids, increased the number of motor units that our decomposition algorithm could theoretically discriminate, i.e., up to 83.5% of the simulated pool (range across conditions: 30.5-83.5%). We then identified motor units from experimental EMG signals recorded in six participants with grids of various sizes (range: 2-36 cm^2^) and IED (range: 4-16 mm). The configuration with the largest number of electrodes and the shortest IED maximized the number of identified motor units (56±14; range: 39-79) and the percentage of early recruited motor units within these samples (29±14%). Finally, the number of identified motor units further increased with a prototyped grid of 256 electrodes and an IED of 2 mm. Taken together, our results showed that larger and denser surface grids of electrodes allow to identify a more representative pool of motor units than currently reported in experimental studies.

**Significance statement:** The application of source separation methods to multi-channel EMG signals recorded with grids of electrodes enables users to accurately identify the activity of individual motor units. However, the design parameters of these grids have never been discussed. They are usually arbitrarily fixed, often based on commercial availability. Here, we showed that using larger and denser grids of electrodes than conventionally proposed drastically increases the number of identified motor units. The samples of identified units are more balanced between early- and late-recruited motor units. Thus, these grids provide a more representative sampling of the active motor unit population. Gathering large datasets of motor units using large and dense grids will impact the study of motor control, neuromuscular modelling, and human-machine interfacing.

## Introduction

Decoding the neural control of natural behaviors relies on the identification of the discharge activity of individual neural cells. Classically, arrays of electrodes are implanted close to the cells to record their electrical activity. The application of algorithms that separate the simultaneous and overlapping activity of these cells has enabled researchers to study neural processes in multiple areas of the brain (Stringer et al., 2019), such as in the motor or the sensorimotor areas (Churchland and Shenoy, 2007; Gallego et al., 2020). At the periphery of the nervous system, it is also possible to record the activity of individual motor neurons innervating muscle fibers (Duchateau and Enoka, 2011; Heckman and Enoka, 2012; Farina et al., 2016). The motor unit, i.e., a motor neuron and the muscle fibers it innervates, acts as an amplifier of the neural activity, as one action potential propagating along a motor neuron’s axon generates an action potential in each of the innervated muscle fibers. The discharge activity of motor units can be identified by decomposing surface electromyographic (EMG) signals into trains of motor unit action potentials (MUAPs) using, e.g., blind-source separation algorithms (Holobar and Farina, 2014; Farina and Holobar, 2016). The multiple observations for source separation are obtained by recording EMG signals with grids of electrodes. This approach usually allows for the reliable analysis of 5 to 40 concurrently active motor units (Del Vecchio et al., 2017; Del Vecchio et al., 2020; Hug et al., 2021a).

While the design of intracortical (e.g., (Jun et al., 2017; Steinmetz et al., 2018)) and intramuscular (e.g., (Muceli et al., 2015; Muceli et al., 2022)) electrodes arrays has scaled up over the years to record larger samples of neural cells, the configuration of grids of surface EMG electrodes has not systematically evolved. Most researchers currently use grids with 64 electrodes arranged in 13 × 5 or 8 × 8 montages, the interelectrode distance (IED) between adjacent electrodes (e.g., 4 mm, 8 mm, or 10 mm) being dictated by the size of the muscle to cover. Yet, optimizing these parameters, i.e., grid size and IED, may influence the performance of EMG decomposition. Currently, there are no recommendations on optimal designs for grids of electrodes.

Source separation algorithms are based on the necessary condition that identifiable motor units have a unique representation of their action potentials across the multi-channel EMG signals (Farina et al., 2008; Holobar and Farina, 2014; Farina and Holobar, 2016). This implies that the three-dimensional waveform of a MUAP (one time dimension and two spatial dimensions) is unique within the pool of active motor units detected by the grid of electrodes. In practice, the identified motor units are those that innervate larger numbers of muscle fibers, as their action potentials tend to have the largest energy. Conversely, low-threshold motor units usually remain hidden since their energy is close to the baseline noise. Increasing the density of electrodes would increase the spatial sampling of EMG signals (Farina and Holobar, 2016), which in turn should improve the discrimination of MUAPs, allowing the identification of a higher number of motor units. Additionally, increasing the density of electrodes may reveal the hidden low-threshold motor units by sampling their action potentials across a higher number of electrodes, leading to a better compensation of the additive noise in the mixture model of the EMG signal (Farina and Holobar, 2016).

In this study, we combined computational and laboratory experiments to identify the optimal design parameters of grids of surface electrodes with the aim to maximize the number of identified motor units. We first simulated a pool of 200 motor units and the associated EMG signals recorded from grids of electrodes of various sizes and densities. These simulations showed that the greater the size and the density of the grid, the higher the percentage of theoretically identifiable motor units and the relative ratio of theoretically identifiable deep units. We confirmed these theoretical results with experimental signals recorded with a grid of 256 electrodes with a 4-mm IED that was down-sampled in the space domain to obtain six grid configurations (surface range: 2-36 cm^2^ and IED range: 4-16 mm). Finally, we prototyped a new grid of 256 electrodes with a 2-mm IED and demonstrated that the number of identified motor units further increased with 2-mm IED. The entire dataset (raw and processed data) and codes are available at https://figshare.com/s/f4a94d9bdff470bf10f8.

## Methods

### Computational study

A pool of 200 motor units was simulated to test whether increasing the density and the size of surface grids of electrodes would impact the number of theoretically identifiable motor units. The simulations were based on an anatomical model entailing a cylindrical muscle volume with parallel fibers (Farina et al., 2008; Konstantin et al., 2020), in which subcutaneous and skin layers separate the muscle from the surface electrodes. Specifically, we set the radius of the muscle to 25.4 mm and the thicknesses of the subcutaneous and skin layers to 5 mm and 1 mm, respectively. The centers of the motor units were distributed within the cross section of the muscle using a farthest point sampling technique. The farthest point sampling filled the cross-section by iteratively adding centers points that were maximally distant from all the previously generated motor unit centers, resulting in a random and even distribution of the motor unit territories within the muscle. The number of fibers innervated by each motor neuron followed an exponential distribution, ranging from 15 to 1500. The fibers of the same motor unit were positioned around the center of the motor unit within a radius of 0.2 to 9.8 mm, and a density of 20 fibers/mm^2^. Because motor unit territories were intermingled, the density of fibers in the muscle reached 200 fibers/mm^2^. The MUAPs were detected by circular surface electrodes with a diameter of 1 mm. The simulated grids were centered over the muscle in the transverse direction, with a size ranging from 14.4 to 36 cm^2^, and an IED ranging from 2 to 36 mm.

### Laboratory study

#### Participants

Six healthy participants (all males; age: 26 ± 4 yr; height: 174 ± 7 cm; body weight: 66 ± 15 kg) volunteered to participate in the first experimental session of the study. They had no history of lower limb injury or pain during the months preceding the experiments. One of these individuals (age: 26 yr; height: 168 cm; bodyweight: 51 kg) participated in a second experimental session to test the prototyped grid with an IED of 2 mm. The Ethics Committee at Imperial College London reviewed and approved all procedures and protocols (no. 18IC4685). All participants provided their written informed consent before the beginning of the experiment.

#### Experimental tasks

The two experimental sessions consisted of a series of isometric ankle dorsiflexions performed at 30% and 50% of the maximal voluntary torque (MVC) during which we recorded high density electromyographic (HD-EMG) signals over the Tibialis Anterior muscle (TA). The participant sat on a massage table with the hips flexed at 30°, 0° being the hip neutral position, and their knees fully extended. We fixed the foot of the dominant leg (right in all participants) onto the pedal of a commercial dynamometer (OT Bioelettronica, Turin, Italy) positioned at 30° in the plantarflexion direction, 0° being the foot perpendicular to the shank. The thigh was fixed to the massage table with an inextensible 3-cm-wide Velcro strap. The foot was fixed to the pedal with inextensible straps positioned around the proximal phalanx, metatarsal and cuneiform. Force signals were recorded with a load cell (CCT Transducer s.a.s, Turin, Italy) connected in-series to the pedal using the same acquisition system as for the HD-EMG recordings (EMG-Quattrocento; OT Bioelettronica). The dynamometer was positioned accordingly to the participant’s lower limb length and secured to the massage table to avoid any motion during the contractions.

All experiments began with a warm-up, consisting of brief and sustained ankle dorsiflexion performed at 50% to 80% of the participant’s subjective MVC. During the warm-up, all participants learnt to produce isometric ankle dorsiflexion without co-contracting the other muscles crossing the hip and knee joints. At the same time, we iteratively adjusted the tightening and the position of the straps to maximize the comfort of the participant. Then, each participant performed two 3-to-5 s MVC with 120 s of rest in between. The peak force value was calculated using a 250-ms moving average window, and then used to set the target level during the submaximal contractions. After 120 s of rest, each participant performed two trapezoidal contractions at 30% and 50% MVC with 120 s of rest in between, consisting of linear ramps up and down performed at 5%/s and a plateau maintained for 20 s and 15 s at 30% and 50% MVC, respectively. The order of the contractions was randomized. One participant (S2) did not perform the contractions at 50% MVC.

#### High-density electromyography

In the first experimental session, four adhesive grids of 64 electrodes (13 x 5 with a missing electrode in a corner; gold coated; 1 mm diameter; 4 mm IED; OT Bioelettronica) were placed over the belly of the TA. The grids were carefully positioned side-to-side with a 4-mm-distance between the electrodes at the edges of adjacent grids (Figure 1A). The 256 electrodes were centered to the muscle belly and laid within the muscle perimeter identified through palpation. The skin was shaved, abrased and cleansed with 70% ethyl alcohol. Electrode-to-skin contact was maintained with a bi-adhesive perforated foam filled with conductive paste. The grids were wrapped with tape and elastic bands to secure the contact with the skin. The four pre-amplifiers were connected in-series with stackable cables to a wet reference band placed above the medial malleolus of the same leg. HD-EMG signals were recorded in monopolar derivation with a sampling frequency of 2,048 Hz, amplified (x150), band-pass filtered (10–500 Hz), and digitized using a 400 channels acquisition system with a 16-bit resolution (EMG-Quattrocento; OT Bioelettronica).

**Figure 1:**
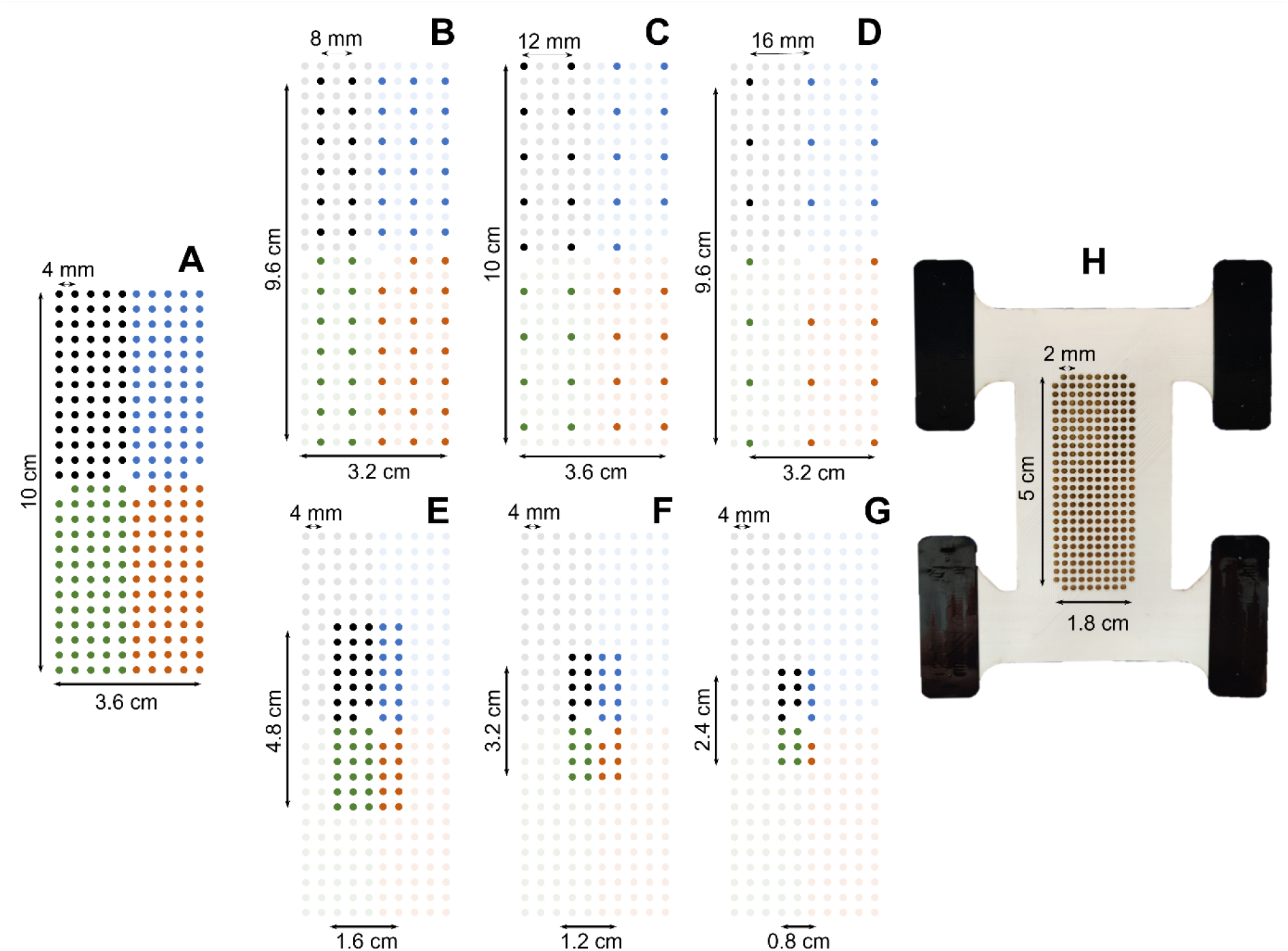
The eight grid configurations considered in this study. From the first grid of 256 electrodes (A, grid size: 36 cm^2^, IED: 4 mm), six shallower and smaller grids (B-G) were artificially obtained by discarding the relevant electrodes. (B,C,D) Density analysis: 8, 12, and 16mm IED. (E,F,G) Size analysis: 7.7, 3.6, and 2 cm^2^ surface area. (H) The ultra-dense prototyped grid of 256 electrodes (grid size: 9 cm^2^, IED: 2 mm).

In the second experimental session, one ultra-dense prototyped grid of 256 electrodes (Figure 1H; 26 x 10 with a missing electrode in each corner; gold coated; 1 mm diameter; 9 cm^2^ area; 2-mm IED; custom-manufactured for this study by OT Bioelettronica) was placed over the belly of the TA and the HD-EMG signals were recorded using the same procedure as previously described.

#### Grid configurations

During the first experimental session, we recorded EMG signals from the TA with a total of 256 electrodes covering an area of 36 cm^2^ over the muscle (10 cm x 3.6 cm, 4-mm IED, Figure 1A). To investigate the effect of electrode density, we down-sampled the grid of 256 electrodes by successively discarding rows and columns of electrodes and artificially generating three new grids covering the same area with IEDs of 8 mm, 12 mm, and 16 mm, that involved 256, 64, 35, and 20 electrodes, respectively (Figure 1B-D). It is noteworthy that the 8-mm and 16-mm grids covered a surface of 32 cm^2^ because they included an odd number of rows and columns. To investigate the effect of the size of the grid, we discarded the peripherical electrodes to generate grids of 63, 34 and 19 electrodes with a 4-mm IED, covering areas of 7.7, 3.8 and 2 cm^2^ over the muscle (Figure 1E-G). We chose these sizes to match the number of electrodes used in the density analysis, thus comparing grids with similar number of electrodes, but different densities and sizes (in Figure 1, B versus E, and C versus F).

During the second experimental session, we recorded EMG signals from the TA with an ultra-dense grid of 256 electrodes covering an area of 9 cm^2^ over the muscle (5 cm x 1.8 cm, 2-mm IED, Figure 1H). Using the same procedure as above, we generated two artificial grids of 64 and 32 electrodes with an IED of 4 mm and 8 mm, respectively.

#### HD-EMG decomposition

We decomposed the signals recorded in all the conditions using the same algorithm, parameters, and procedure. First, the monopolar EMG signals were band-pass filtered between 20 and 500 Hz with a second-order Butterworth filter. The channels with low signal-to-noise ratio or artifacts were discarded after visual inspection. The HD-EMG signals were then decomposed into individual motor unit pulse trains using convolutive blind-source separation, as previously described (Negro et al., 2016). In short, the EMG signals were first extended by adding delayed versions of each channel. We kept the same extension factor for all the conditions to reach 1000 extended channels, as previously suggested (Negro et al., 2016). The extended signals were spatially whitened to make them uncorrelated and of equal power. Thereafter, a fixed-point algorithm was applied to identify the sources embedded in the EMG signals, i.e., the motor unit pulse trains, or series of delta functions centered at the motor unit discharge times. In this algorithm, the contrast function g(x) = log(cosh(x)) was iteratively applied to the EMG signals to skew the distribution of the values of the motor unit pulse trains toward 0, and thus maximize the level of sparsity of the motor unit pulse train. The high level of sparsity matches the physiological properties of motor units, with a relatively small number of discharges per second (< 50 discharge times/s during submaximal isometric contractions). The convergence was reached once the level of sparsity did not substantially vary (with a tolerance fixed at 10^-4^) when compared to the previous iteration (Negro et al., 2016). At this stage, the motor unit pulse train contained high peaks (i.e., the delta functions from the identified motor unit) and lower values due to the activities of other motor units and noise. High peaks were separated from lower values using peak detection and K-mean classification with two classes. The peaks from the class with the highest centroid were considered as the discharge times of the identified motor unit. A second algorithm refined the estimation of the discharge times by iteratively recalculating the motor unit filter and repeating the steps with peak detection and K-mean classification until the coefficient of variation of the inter-spike intervals was minimized. This decomposition procedure has been previously validated using experimental and simulated signals (Negro et al., 2016). After the automatic identification of the motor units, duplicates were automatically removed. For this purpose, the pulse trains identified from pairs of motor units were first aligned using a cross-correlation function to account for a potential delay due to the propagation time of action potentials along the fibers. Then, two discharge times were considered as common when they occurred within a time interval of 0.5 ms, and two or more motor units were considered as duplicates when they had at least 30% of their identified discharge times in common (Holobar et al., 2010). In principle, the limited level of synchronization between individual motor units results in a few simultaneous discharges between pairs of motor units. A threshold of 30% is therefore highly conservative to ensure the removal of all motor units with a level of synchronization well above physiological values. It is worth noting that most of the motor units identified as duplicates after the automatic decomposition had almost 100% of their discharge times in common. In that case, the motor unit with the lowest coefficient of variation of the inter-spike intervals was retained for the analyses. At the end of these automatic steps, all the motor unit pulse trains, i.e., the output of the decomposition resulting from the projection of EMG signals onto individual motor unit filters, were visually inspected, and manual editing was performed to correct the false identification of artifacts or the missed discharge times (Del Vecchio et al., 2020; Hug et al., 2021b; Avrillon et al., 2023). The update of the motor unit filters with the corrected discharge times and the recalculation of the motor unit pulse trains always improved the distance between the discharge times and the noise, quantified with the pulse-to-noise ratio (PNR) (Holobar et al., 2014). Note that this manual step is highly reliable across operators, as previously demonstrated by Hug et al. (2021b). Duplicates were checked a second time after manual editing, with very rare cases of removal as most of the duplicates were automatically identified after the automatic decomposition. Only the motor unit pulse trains which exhibited a PNR > 28 dB after manual editing were retained for further analysis.

We further tested whether decomposing subsets of electrodes within a highly populated grid of 256 electrodes increased the number of identified motor units. Indeed, the lower ratio of large motor units sampled by each independent subset of 64 electrodes could allow the algorithm to converge to smaller motor units that contribute to the signal. For a similar number of iterations, it is likely that these motor units would have otherwise contributed to the noise component of the mixture model of the EMG signal (Farina and Holobar, 2016). Thus, we decomposed the grids of 256 electrodes (4-mm and 2-mm IED, Figure 1A, H) as four separated grids of 64 electrodes before removing the motor units duplicated between grids.

### Analyses

#### Computational study

We first estimated the theoretical percentage of identifiable motor units for each of the simulated conditions. To do so, the simulated MUAPs detected over the entire set of electrodes were compared with each other. The comparisons were done pairwise by first aligning the MUAPs in time using the cross-correlation function, and then computing the normalized mean square difference between the aligned action potentials. Pairs of action potentials with a mean square difference below 5% were considered not discriminable. The 5% criterion was based on the variability of motor unit action potential shapes observed experimentally for individual motor units (Farina et al., 2008). After computing all pair-wise comparisons, we then computed the percentage of action potentials that could be discriminated from all others, i.e., the theoretical percentage of identifiable motor units. This metric is independent from the algorithm used for decomposition and establishes a theoretical upper bound in the number of motor units that can be identified by any decomposition algorithm. For each theoretically identifiable motor unit, we also computed the distance between the center of the territory of the corresponding muscle fibers and the skin surface.

#### Laboratory study – number of identified motor units

We reported the absolute number of motor units (PNR > 28 dB) identified with all the grid configurations. For each participant, the number of identified motor units was then normalized to the maximal number of motor units found across all conditions, yielding normalized numbers of identified motor units 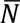 expressed in percentage. For each condition, we calculated the mean and standard deviation of the 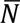 values across participants. To investigate the effects of density and size of the grid, we fitted logarithmic trendlines to the relationships between the averaged 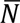 values and IED or grid size. We also fitted a logarithmic trendline to the average 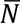values and their corresponding number of electrodes, in which case the conditions involving the same number of electrodes, but different grid size and density, were given a weight of 0.5 in the minimization function. We reported the r^2^ and p-value for each regression trendline. To maintain consistency with the computational study investigating the number of theoretically identifiable motor units across grid designs, the trendlines were fitted on the results obtained when the complete grids of 256 electrodes were decomposed as independent subsets of 64 electrodes, which systematically returned the highest number of identified motor units. The trendlines fitted on the results obtained with the decomposition of the 256 electrodes as a whole are reported in Figure 4-1.

#### Laboratory study – properties of identified motor units

To investigate the effects of electrode density and grid size on the properties of the identified motor unit, we used a typical frequency distribution of the motor unit force recruitment thresholds in the human TA (Caillet et al., 2022b), where *F*^*th*^(*j*) is the force recruitment threshold of the j^th^ motor unit in the normalized motor unit pool ranked in ascending order of *F*^*th*^.

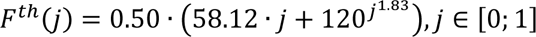

The identified motor units were then classified according to this relationship and their measured force recruitment threshold, the first half of the active pool being ‘early recruited’, and the second half ‘late recruited’ (Henneman and Mendell, 1981; Caillet et al., 2022a). For each condition, we reported the percentage of identified motor units that were ‘early recruited’. We did not report this metric when five or fewer motor units were identified in one condition for three or more participants.

#### Laboratory study – correlation between observations

We assessed how the density of electrodes impacted the information redundancy in EMG signals recorded by adjacent electrodes. To this end, MUAP shapes were identified over the 256 electrodes with the spike-triggered averaging technique. To do so, the discharge times were used as a trigger to segment and average the HD-EMG signals over a window of 50 ms. For each motor unit, we identified the electrode with the highest action potential peak-to-peak amplitude and calculated the average correlation coefficient ρ between this action potential and those recorded by the four adjacent electrodes with an IED of 4 mm, 8 mm, 12 mm, and 16 mm. We also repeated this correlation analysis for the ultra-dense grid of 256 electrodes using an IED of 2 mm, 4 mm, and 8 mm.

## Results

All the datasets (raw and processed data) and codes used to process the data are available at https://figshare.com/s/f4a94d9bdff470bf10f8.

### Computational study

We simulated the discharge activity of 200 motor units recorded by 84 configurations of grids of electrodes (Figure 2; surface range: 14.4 to 36 cm^2^, IED range: 2 to 36 mm). The number of theoretically identifiable motor units increased with the size of the grid, from 46.7 ± 7.7% of the motor units theoretically identifiable with a grid of 14.4 cm^2^ to 77.8 ± 5.5% of the motor units theoretically identifiable with a grid of 36 cm^2^. The number of theoretically identifiable motor units also increased with shorter interelectrode distances. For example, with a grid of 36 cm^2^, the number of theoretically identifiable motor units increased from 63.5% to 83.5% of the motor units with an IED of 36 and 2 mm, respectively (Figure 2B). Increasing the surface size and the density of the grid of electrodes revealed deeper motor units. The averaged distance of theoretically identifiable motor units from the skin increased with the size of the grid (Figure 2C; 14.3 ± 0.1 mm vs. 16.5 ± 0.2 mm with grids of 14.4 and 36 mm^2^, respectively), but not with the IED of the grid (Figure 2D; 15.6 ± 1.1 mm vs. 15.5 ± 0.9 mm with an IED of 36 and 2 mm, respectively).

**Figure 2:**
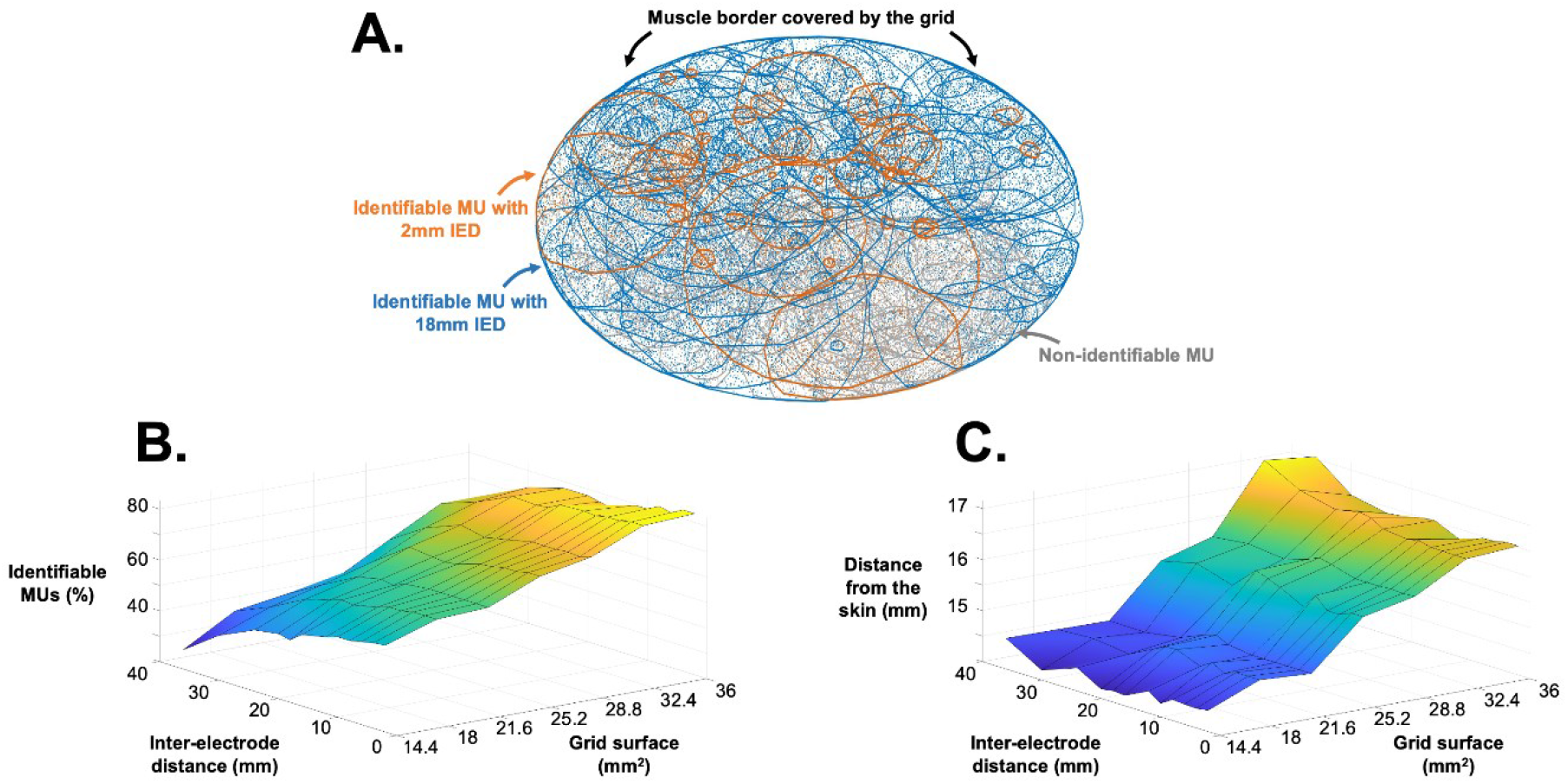
Results from the 200 simulated motor units with 84 configurations of grids of electrodes. (A) Each solid line represents a motor unit territory, the scatters being the muscle fibers. Blues lines are the theoretically identifiable motor units with a grid of 21.6 cm^2^ and an interelectrode distance (IED) of 18 mm, while the orange lines are the motor units revealed with a grid of 21.6 cm^2^ and an IED of 2mm. Grey lines represent the non-identifiable motor units. The percentage of theoretically identifiable motor units (B) and their distance from the skin (C) are reported for the 84 configurations.

### Laboratory study - grids of 256 electrodes with an IED of 4-mm

#### Number of identified motor units

The motor unit pulse trains automatically identified across all conditions, intensities, and participants were visually inspected and carefully edited when a missing discharge time or a falsely identified artifact were observed. On average, 9 ± 4 % and 22 ± 9 % of the motor units automatically identified at 30% and 50% MVC, respectively, were removed after visual inspection and manual editing. Furthermore, when the four grids of 64 electrodes were separately decomposed, 30 ± 5 % and 24 ± 6 % of the automatically identified motor units were removed because they were identified in more than one grid (only one pulse train was retained in case of duplicates). The highest number of identified motor units was systematically reached with the separate decomposition of the four grids of 64 electrodes with an IED of 4 mm, with 56 ± 14 motor units (PNR = 34.2 ± 1.1) and 45 ± 10 motor units (PNR = 34.0 ± 0.9) at 30% and 50% MVC, respectively (Figure 3). At least 82% of the motor units identified in one condition were also identified in the conditions involving a higher number of electrodes. Similarly, 91% to 100% of the motor units identified in one condition were also identified with the 256-electrode configuration (4-mm IED, 36-cm^2^ size, Figure 1A) with the four grids decomposed separately.

**Figure 3:**
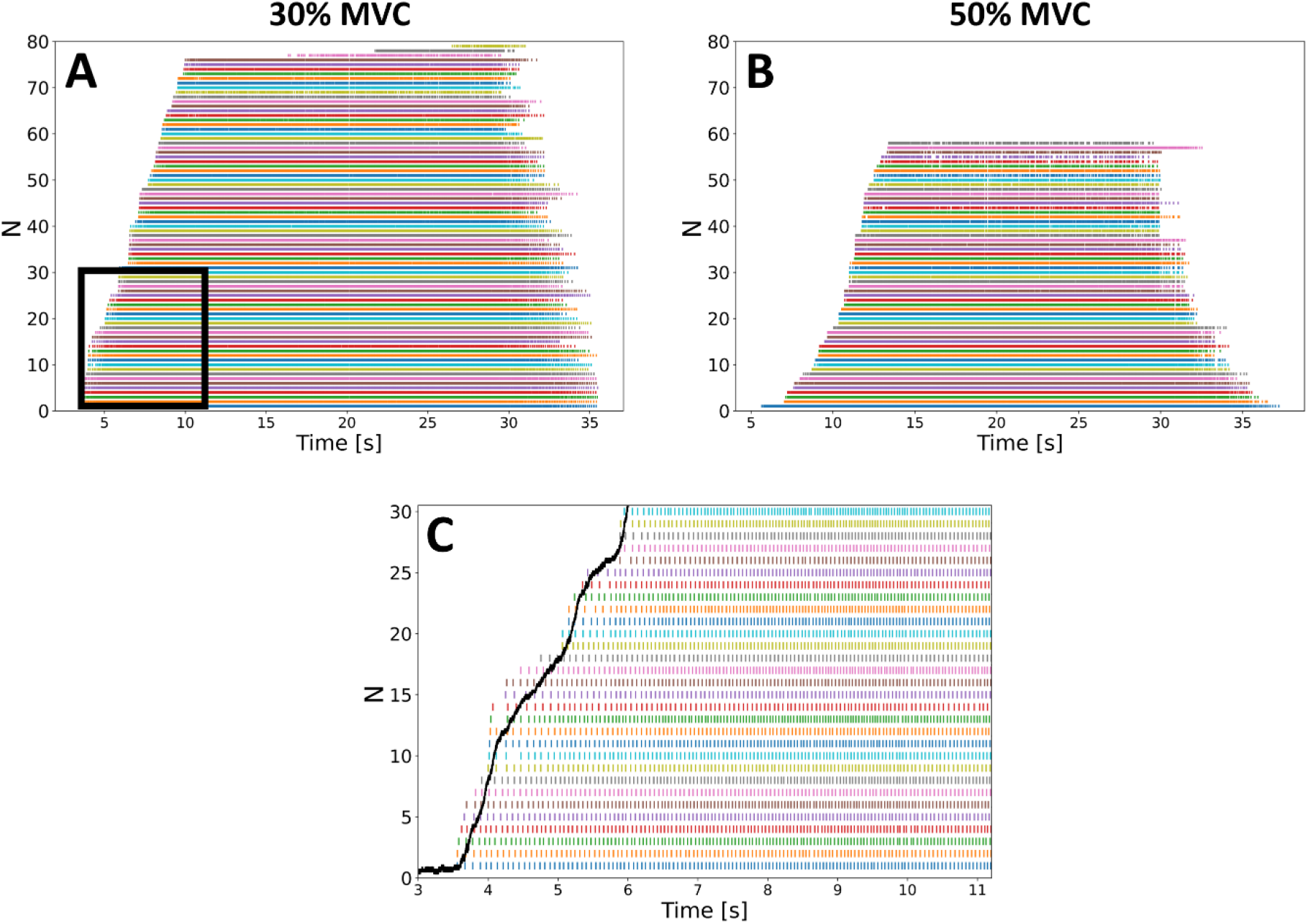
Discharge times of the maximum number of motor units identified in one participant (S1) at 30% (A) and 50% MVC (B), with 79 and 58 identified motor units, respectively. The motor units were identified with separated decompositions of the four grids of 64 electrodes (4 mm IED). (C) Discharge times of the 30 first recruited motor units during the ascending ramp of force (black curve) at 30% MVC (black box in A).

When considering the effect of electrode density (grid size fixed at 32-36 cm^2^, Figure 1A-D), we found the lowest number *N* of identified motor units with the 16-mm IED, with 3 ± 1 motor units and 2 ± 1 motor units at 30% and 50% MVC, respectively (Figure 4A, C). Additional motor units were gradually identified with greater electrode densities. The highest number of identified motor units was observed with the highest density (4-mm IED), with 56 ± 14 and 45 ± 10 motor units at 30% and 50% MVC, respectively, with the 4×64-electrode decomposition procedure (Figure 4A, C). With the 256-electrode decomposition procedure, 43 ± 11 and 25 ± 6 motor units were identified at 30% and 50% MVC, respectively (Figure 4A, C). Finally, we found a decreasing logarithmic relationship between the normalized number 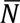 of motor units, averaged for each participant, and the IED, with r^2^ = 1.0 (p = 2.5·10^-5^) and r^2^ = 0.99 (p = 0.001) at 30% and 50% MVC, respectively (Figure 4B, D).

**Figure 4:**
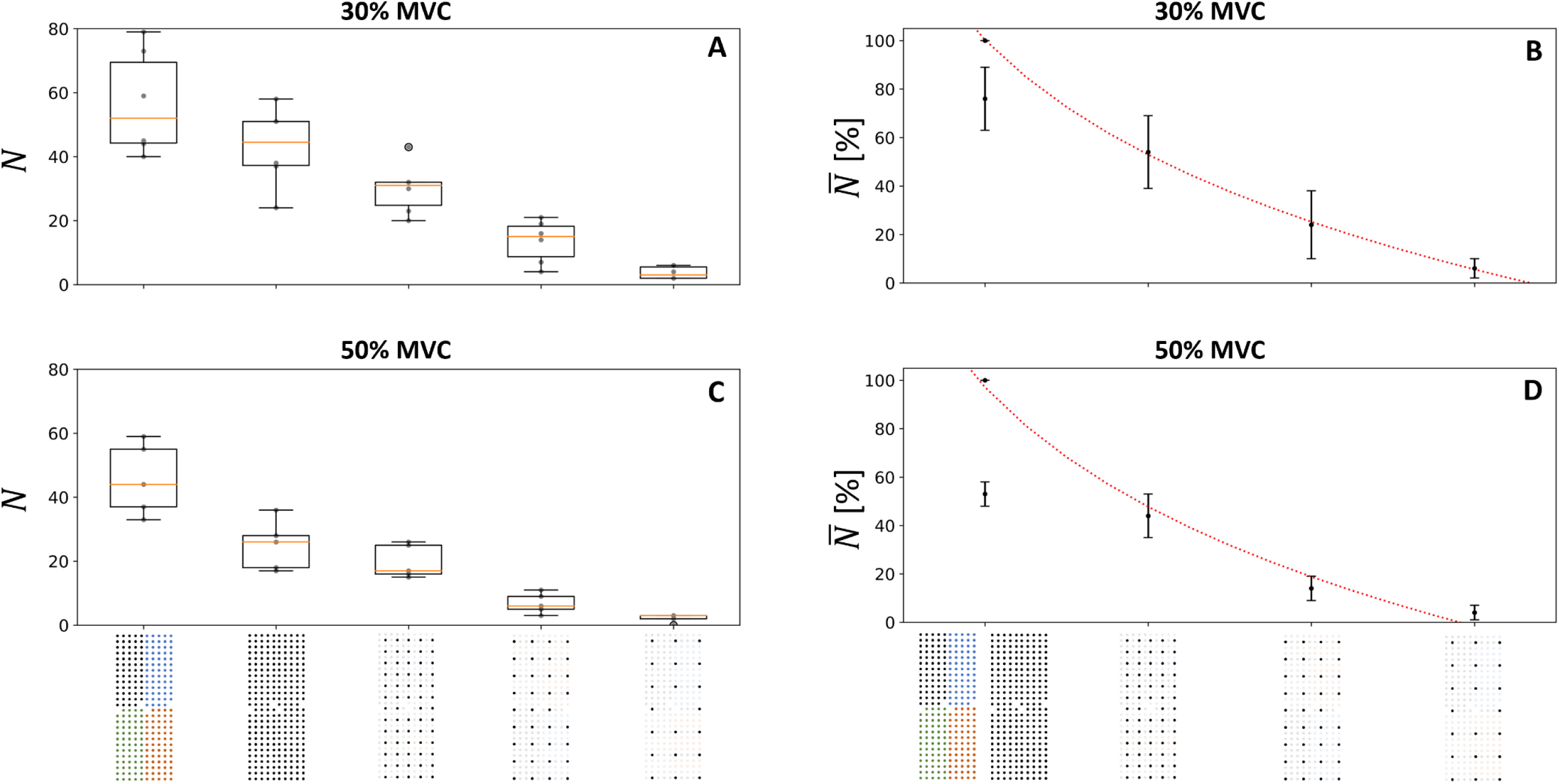
Effect of the electrode density on the number of identified motor units *N* at 30% (A, B) and 50% MVC (C, D). The boxplots in the left column report the absolute er *N* of identified motor units per participant (grey dots) and the median (orange line), quartiles, and 95%-range across participants. In the right column, the normalized er of motor units 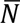 ogarithmically decreases with interelectrode distance *d* (4, 8, 12, and 16mm in abscissa) as 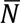 = 195 − 68 *log*(*d*) (*r*^2^ = 1.0, *p* = 2.5 · 10^−5^) at 30% (B) and 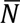 = 196 − 71 *log*(*d*) (*r*^2^ = 0.99, *p* = 0.001) at 50% MVC (D). The standard deviation of 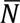 across subjects is displayed with vertical bars. Moreover, the y of the motor unit pulse trains (i.e., decomposition accuracy, estimated by the PNR) increased when increasing the density of electrodes (see Figure 4-3 for more details). ecomposition procedures were considered for the 256-electrode condition; the grid of 256 black electrodes indicates that the 256 signals were simultaneously decomposed e grid of 256 electrodes of four different colors indicates that four subsets of 64 electrodes were decomposed. To maintain consistency with the computational study, the ines were fitted with the 4*64 condition, which returned the higher number of identified motor units (see Figure 4-1 for the other fitting condition). It is worth noting that utationally increasing the density of electrodes by resampling the EMG signals with a spatial interpolation did not reveal any previously hidden motor units (Figure 4-2).

**Figure 4-1.**
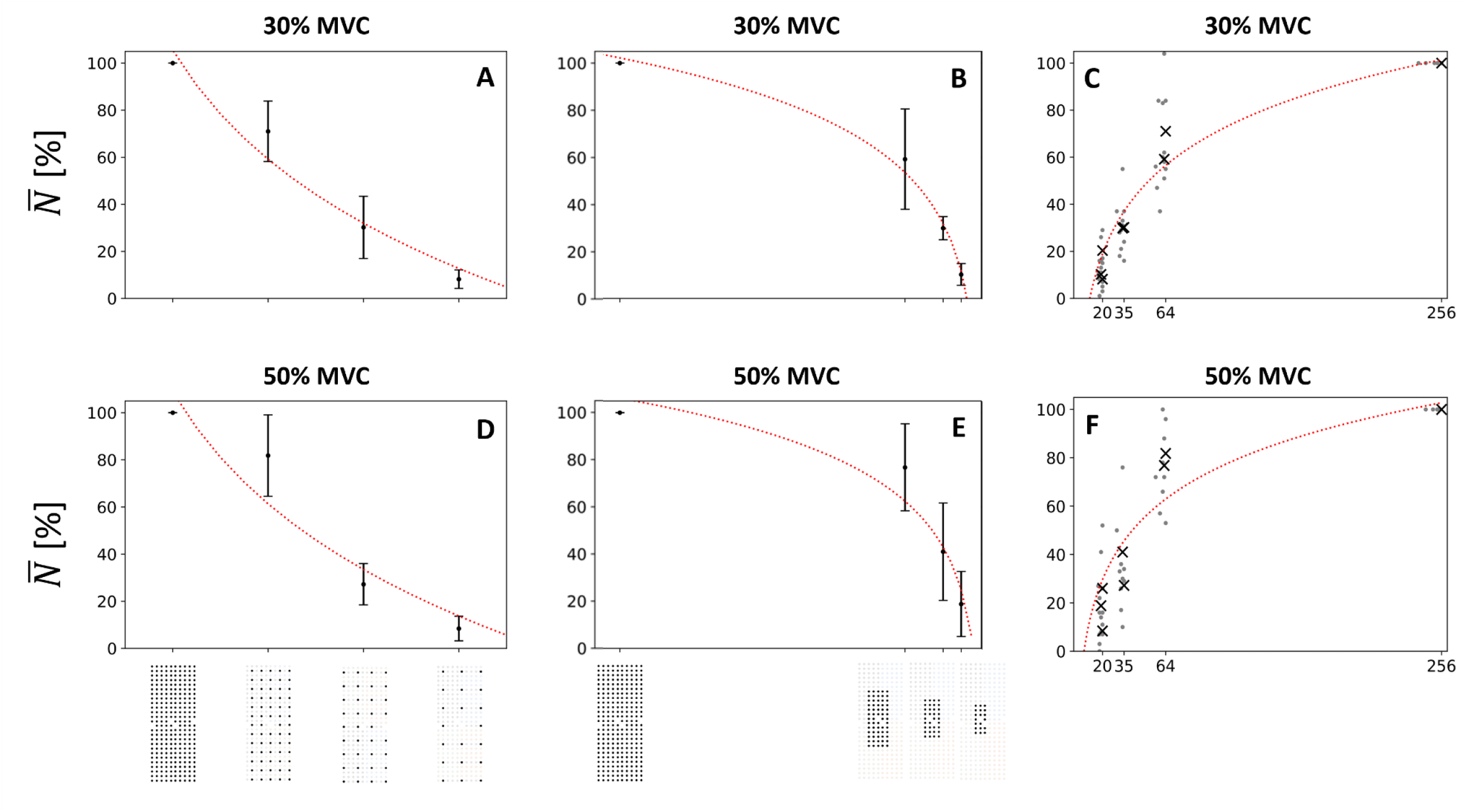
Effect of the density of the grid (A, D), the size of the grid (B, D), and the number of electrodes (C, F) on the normalized number 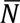 of identified motor units at 30% C) and 50% MVC (D, E, F). 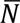 was estimated after decomposing the full grid of 256 electrodes and manually editing the motor unit pulse trains. Vertical bars (A, B, D, the standard deviation of 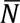 across subjects, scatters are the individual data points, and crosses are their mean (C, F). Logarithmic trendlines were fitted between the ed values 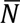 and IED, grid size, and number of channels, as in Figures 4, 5, and 6 of the main document. Here, the trendlines were fitted with the values obtained from composition of the full grid of 256 electrodes. Consistent with the results provided in the main document, 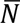 increased with electrode density (*d*), grid size (*s*), and with umber of electrodes (*n*) following statistically significant logarithmic trendlines (p < 0.05). At 30% MVC, 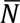 = 198 − 67 *log*(*d*) (*r*^2^ = 0.92), 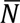 = −10 + (*s*) (*r*^2^ = 0.98), and 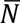 = −78 + 32 *log*(*n*) (*r*^2^ = 0.90). At 50% MVC, 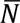 = 204 − 69 *log*(*d*) (*r*^2^ = 0.92), 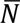 = 5 + 28 *log*(*s*) (*r*^2^ = 0.98), and 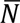 = −57 + (*n*) (*r*^2^ = 0.90). It is noteworthy that the trendlines exhibited more pronounced plateaus (lower *b* value in the *y* = *a* + *b* · log(*x*) trendlines) with the decomposition of l grid of 256 electrodes than with the decomposition of subsets of 64 electrodes.

**Figure 4-2.**
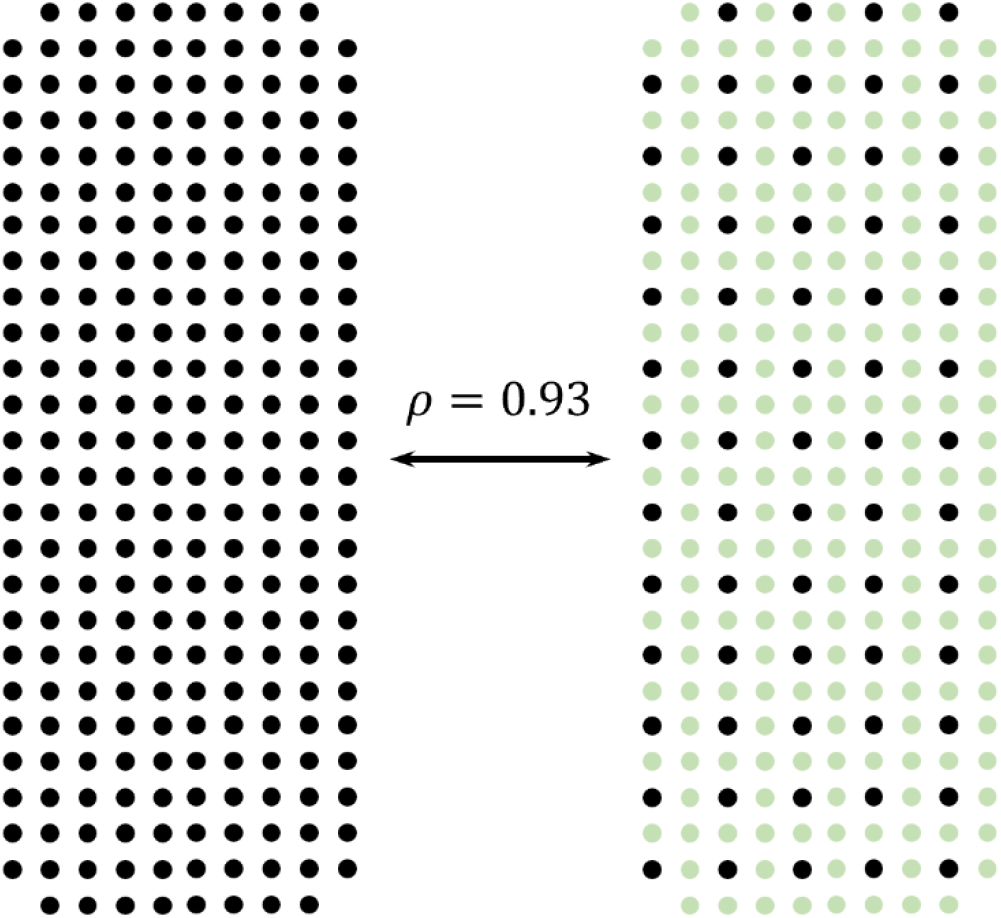
Correlation ρ between experimentally recorded (Left, black) and interpolated (Right, green) EMG signals (Right, black). Using the ultra-dense grid of 256 electrodes (2-mm IED) at 30% MVC, we spatially interpolated down-sampled montages of 4×9 electrodes with an IED of 8 mm and 5×13 electrodes with an IED of 4 mm to generate 5×13 (4-mm IED) and 10×26 (2-mm IED) grids of electrodes, respectively. In these interpolated grids, 25% of the signals were therefore experimentally recorded (Right, black) and 75% interpolated (Right, green). After comparing interpolated and experimentally recorded grids of electrodes, we observed that a better signal reconstruction was obtained with the 2-mm IED, with a correlation coefficient of ρ = 0.93 ± 0.09 between recorded and interpolated signals. We identified 4 and 19 motor units from the interpolated grid with a 4-mm and 2-mm IED, respectively, vs. 19 and 24 motor units with the experimentally recorded signals. We only identified the same motor units as identified with the original less dense grids used to generate the interpolation. These results indicate that interpolation is not sufficient to reconstruct signals from a lower spatial sampling. This may be due to the spatial bandwidth which is greater than the inverse of the minimal interelectrode distance used or to the edge effects of the interpolation due to the relatively small size of the grid.

**Figure 4-3.**
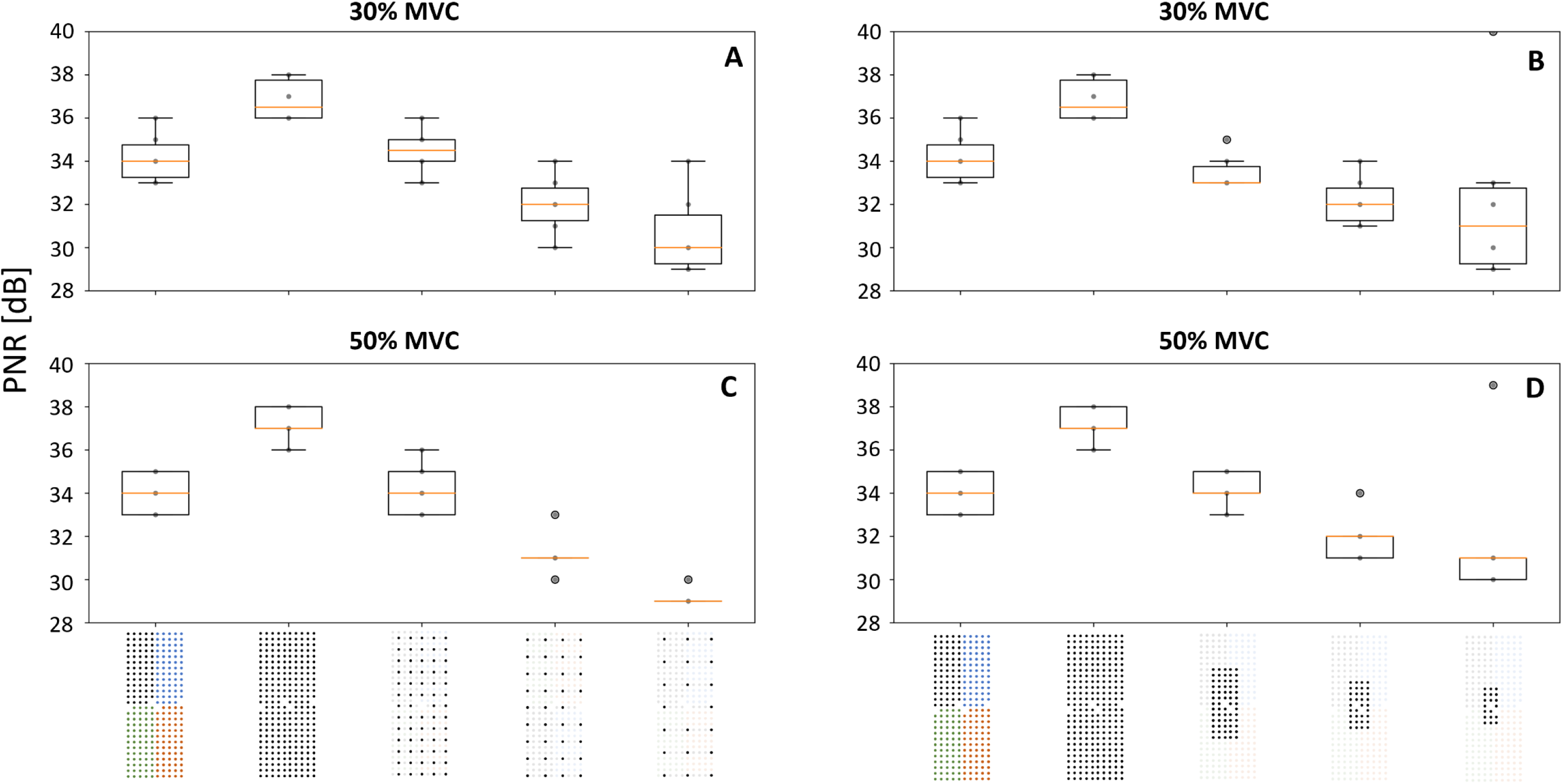
Effect of the electrode density (A, C) and grid size (B, D) on the average PNR across the identified spike trains at 30% MVC (A, B) and 50% MVC (C, D). The boxplots report the average PNRs per participant (grey dots) and the median (orange line), quartiles, and 95%-range across participants. We calculated the average PNR value for the motor unit spike trains (PNR > 28 dB) identified in each subject and condition. The average PNR across identified motor units increased together with both the density and the size of the grid. The lowest PNR values were observed with 16 mm-IED (30 ± 1.8 dB at 30% MVC and 29 ± 1.2 dB at 50% MVC) and with a grid of 2 cm^2^ (31 ± 0.9 dB at 30% MVC and 30 ± 0.9 dB at 50% MVC). The highest PNR was observed with 4 mm-IED and a grid of 36 cm^2^ (36 ± 0.7 dB at 30% MVC and 37 ± 0.7 dB at 50% MVC), enabling the operators to quickly edit the identified motor units.

When considering the effect of the size of the grid (IED fixed at 4 mm, Figure 1A, E-G), we found the lowest number *N* of motor units with a grid of 2 cm^2^, with 4 ± 2 motor units and 4 ± 2 motor units at 30% and 50% MVC, respectively (Figure 6A, C). Additional motor units were then gradually identified with larger grid sizes. The highest number of motor units was observed with a grid of 36 cm^2^, with 56 ± 14 and 45 ± 10 motor units at 30% and 50% MVC, respectively, with the 4×64-electrode decomposition procedure (Figure 4A, C). With the 256-electrode decomposition procedure, 43 ± 11 and 25 ± 6 motor units were identified at 30% and 50% MVC, respectively (Figure 4A, C). Finally, we found an increasing logarithmic relationship between the normalized number of motor units ^*N*^, averaged for each participant, and the size of the grid, with r^2^ = 0.99 (p = 3.0·10^-4^) and r^2^ = 0.98 (p = 0.001) at 30% and 50% MVC, respectively (Figure 6B, D). It is noteworthy that the parameters of the fits were very similar at 30% and 50% MVC in both analyses.

**Figure 5:**
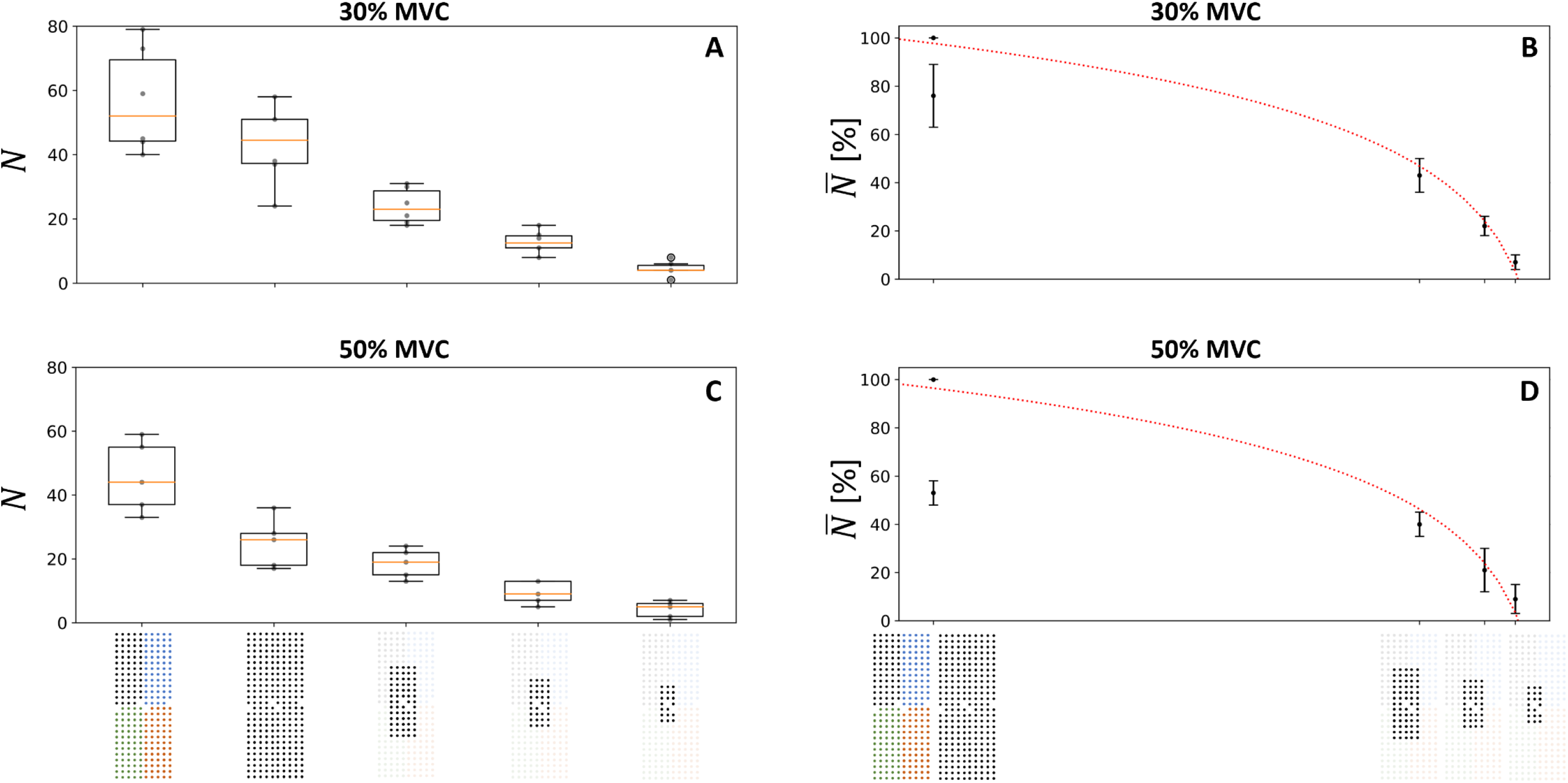
Effect of the size of the grid on the number of identified motor units *N* at 30% (A, B) and 50% MVC (C, D). The boxplots in the left column report the absolute er *N* of identified motor units per participant (grey dots) and the median (orange line), quartiles, and 95%-range across participants. In the right column, the normalized er of motor units 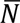 logarithmically decreases with the size of the grid s (2, 3.8, 7.7, and 36 cm^2^ in abscissa) as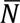 = −20 + 33 *log*(*s*) (*r*^2^ = 0.99, *p* = 3.0 · 10^−4^) at 30% (B), and 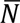 = −19 + 32 *log*(*s*) (*r*^2^ = 0.98, *p* = 0.001) at 50% MVC (D). The standard deviation of 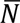 across subjects is displayed with vertical bars. Moreover, the y of the identified motor unit pulse trains (i.e., decomposition accuracy, estimated by the PNR) increased when increasing the size of the grid (see Figure 4-3 for more). Two decomposition procedures were considered for the 256-electrode condition; the grid of 256 black electrodes indicates that the 256 signals were simultaneously posed and the grid of 256 electrodes of four different colors indicates that four subsets of 64 electrodes were decomposed. To maintain consistency with the computational the trendlines were fitted with the 4*64 condition, which returned the higher number of identified motor units (see Figure 4-1 for the other fitting condition).

**Figure 6:**
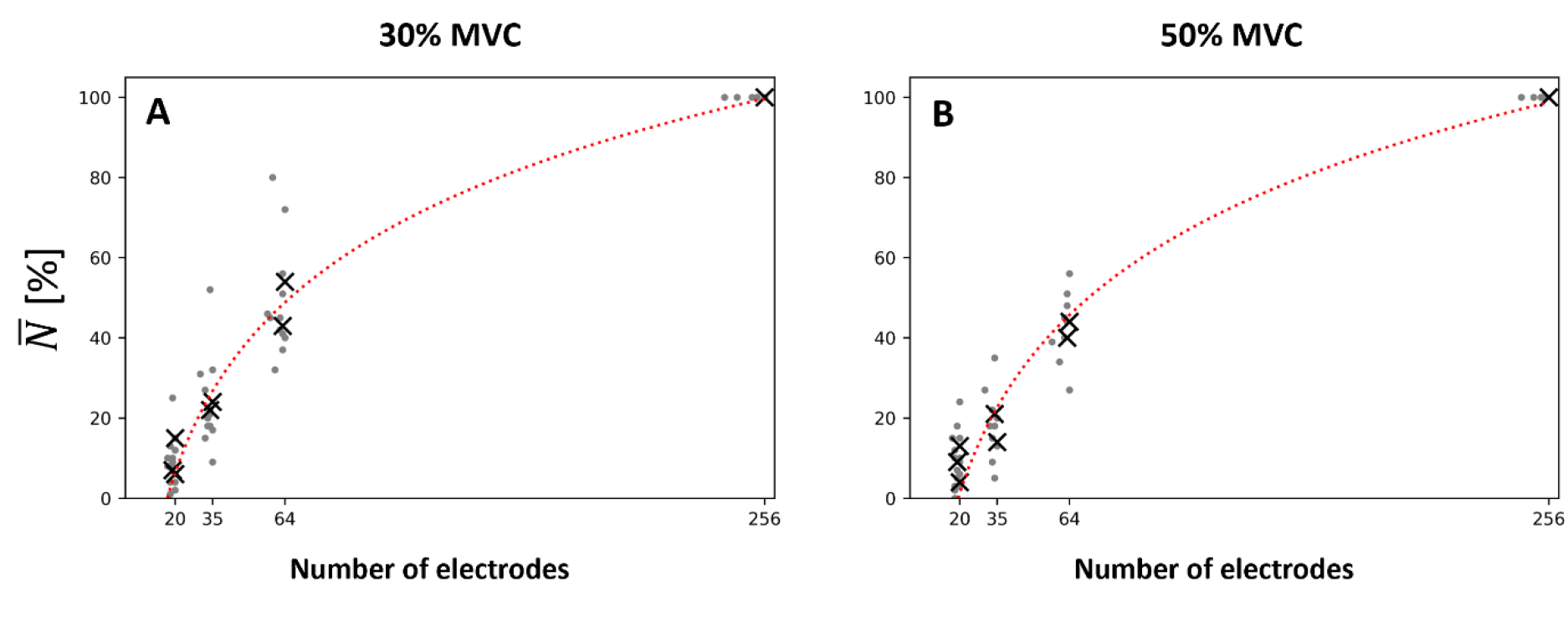
Effect of the number *n* of electrodes on the normalized number 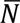 of identified motor units at 30% (A) and 50% MVC (B). The discrete results per participant are displayed with grey data points. The average values 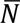 per condition are displayed with black crosses. Weighted logarithmic trendlines were fitted to the data and returned (A) 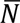 = −104 + 37 *log*(*n*) (*r*^2^ = 0.98, *p* = 0.018), and (B) 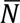 = −113 + 38 *log*(*n*) (*r*^2^ = 0.95, *p* = 0.016). Two decomposition procedures were considered for the 256-electrode condition; the grid of 256 black electrodes indicates that the 256 signals were simultaneously decomposed and the grid of 256 electrodes of four different colors indicates that four subsets of 64 electrodes were decomposed. To maintain consistency with the computational study, the trendlines were fitted with the 4*64 condition, which returned the higher number of identified motor units (see Figure 4-1 for the other fitting condition).

As both the density and the size of the grid determine the number of electrodes, we finally fitted the relationship between the normalized number of motor units 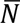 and the number of electrodes. As observed previously, more motor units were identified with a larger number of electrodes, following a logarithmic tendency with r^2^ = 0.98 (p = 0.018) and r^2^ = 0.95 (p = 0.016) at 30% and 50% MVC, respectively (Figure 6). A plateau should theoretically be reached with grids of 1024 and 4096 electrodes (36-cm^2^ grids with 2-mm and 1-mm IED, respectively), with a prediction of 50% and 90% more motor units.

For a fixed number of electrodes, it is noteworthy that the size and the density, although linked, may have different impact on the number of identified motor units (black crosses in Figure 6). For example, 1.25 times more motor units were obtained with the 64-electrode condition (32 cm^2^, 8-mm IED, Figure 1B) than with the 63-electrode condition (7.7 cm^2^, 4-mm IED, Figure 1E) for the group of participants at 30% MVC.

#### Characteristics of identified motor units

We found an increasing logarithmic relationship between the percentage of early recruited motor units for each participant and the density of the grid, with r^2^ = 0.91 (p = 2.8·10^-3^) at 30% MVC (Figure 7F). Contrary to the density, the size of the grid did not impact the percentage of early recruited motor units, with the percentage ranging from 20 to 29% across all sizes, and the logarithmic trendline returning a negligible slope and a low r^2^ = 0.28 (Figure 7C, G). Such differences were also not observed at 50% MVC, where the percentage of early recruited motor units remained below 10% for all conditions.

**Figure 7:**
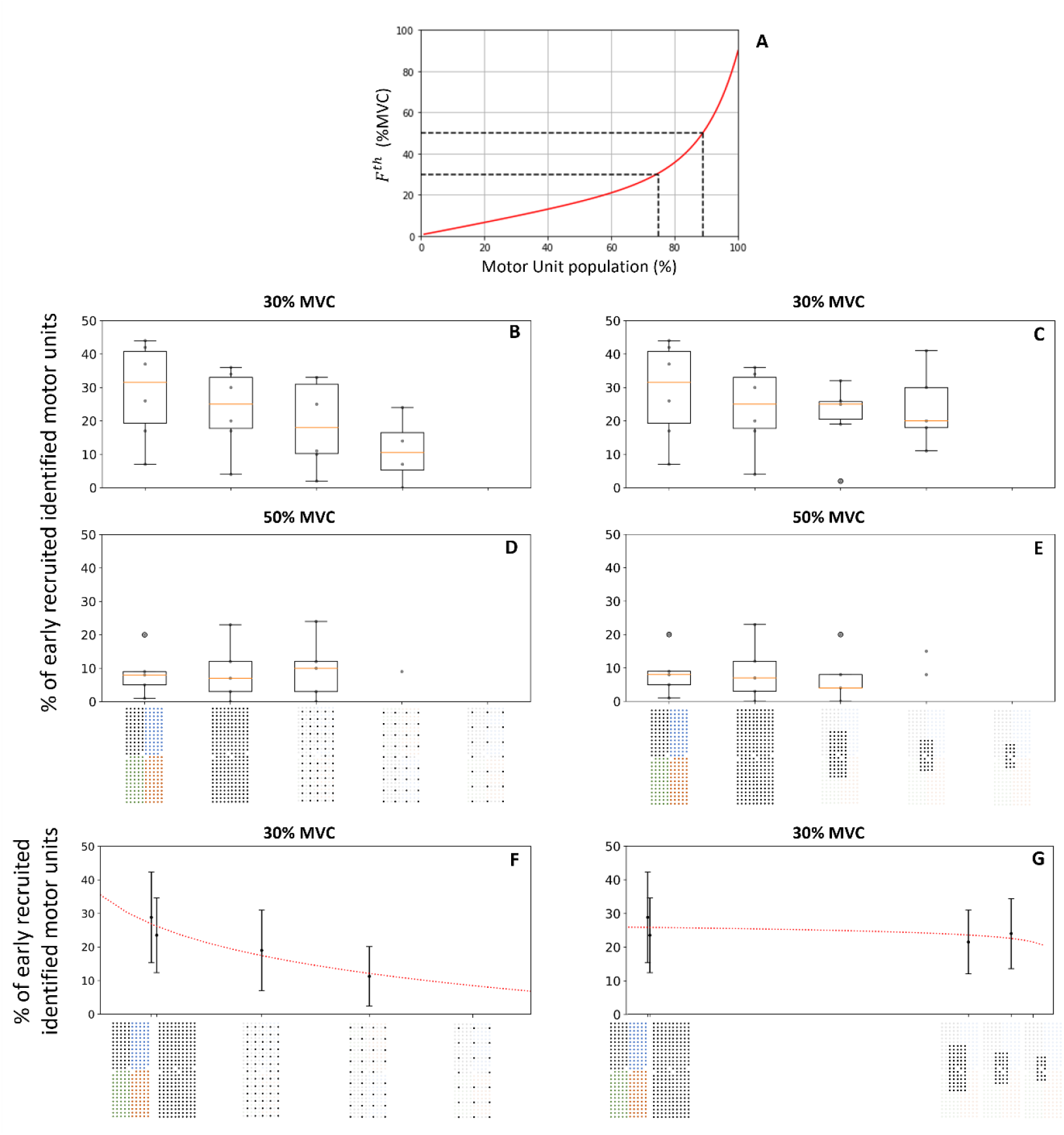
(A) Typical frequency distribution of motor unit force recruitment thresholds in a human TA. The black dashed lines denote the theoretical portions of the population of motor units recruited at 30% and 50% MVC. Effect of the grid density (B, D, F) and grid size (C, E, G) on the percentage of early recruited motor units identified at 30% (B, C, F, G) and 50% MVC (D, E). The boxplots report the results per participant (grey dots) and the median (orange line), quartiles, and 95%-range across participants. (F) At 30% MVC, the percentage of early recruited identified motor units logarithmically decreases with interelectrode distance *d* (4, 8, 12, and 16mm in abscissa) as 44.6 − 13.1 *log*(*d*) (*r*^2^ = 0.91, *p* = 2.8 · 10^−3^). (G) At 30% MVC, the percentage of early recruited identified motor units does not vary with the size of the grid s (2, 3.8, 7.7, and 36 cm^2^ in abscissa), the logarithmic trendline fitting (20.5 + 1.2 *log*(*s*)) returning a negligible slope and low *r*^2^ = 0.28 (*p* = 8 · 10^−4^). The standard deviation across subjects is displayed with vertical bars. Two decomposition procedures were considered for the 256-electrode condition; the grid of 256 black electrodes indicates that the 256 signals were simultaneously decomposed and the grid of 256 electrodes of four different colors indicates that four subsets of 64 electrodes were decomposed. To maintain consistency with the computational study, the trendlines were fitted with the 4*64 condition, which returned the higher number of identified motor units (see Figure 4-1 for the other fitting condition). We did not report the results when five or fewer motor units were identified in one condition for three or more participants.

To support the above observations made at 30% MVC, grids with the same number of electrodes, but different densities and sizes, were directly compared. 62% of the motor units identified with the grids of 64 electrodes (32 cm^2^, IED 8 mm) and 63 electrodes (7.7 cm^2^, IED 4 mm) were identified in both conditions at 30% MVC. 28 ± 9% of the motor units specific to the 8-mm IED grid were early recruited, while 44 ± 11% of the motor units specific to the 4-mm IED condition were early recruited. Similar results were obtained with the grids of 35 (36 cm^2^, 12-mm IED) and 34 electrodes (3.6 cm^2^, 4-mm IED), where a higher number of early recruited motor units were specifically identified with denser rather than larger grids.

#### Correlation between MUAPs from adjacent electrodes

Figure 8 reports the effect of the density of electrodes on the level of correlation ρ between the profiles of action potentials recorded by adjacent electrodes. The lowest average correlation coefficient ρ was observed with an IED of 16 mm (ρ = 0.87 ± 0.03 and ρ = 0.88 ± 0.04 at 30% and 50% MVC, respectively). The level of correlation increased with a shorter IED, with ρ = 0.96 ± 0.04 and ρ = 0.95 ± 0.05 between the profiles of action potentials recorded by adjacent electrodes with a 4-mm IED at 30% and 50% MVC, respectively (Figure 8B, C).

**Figure 8:**
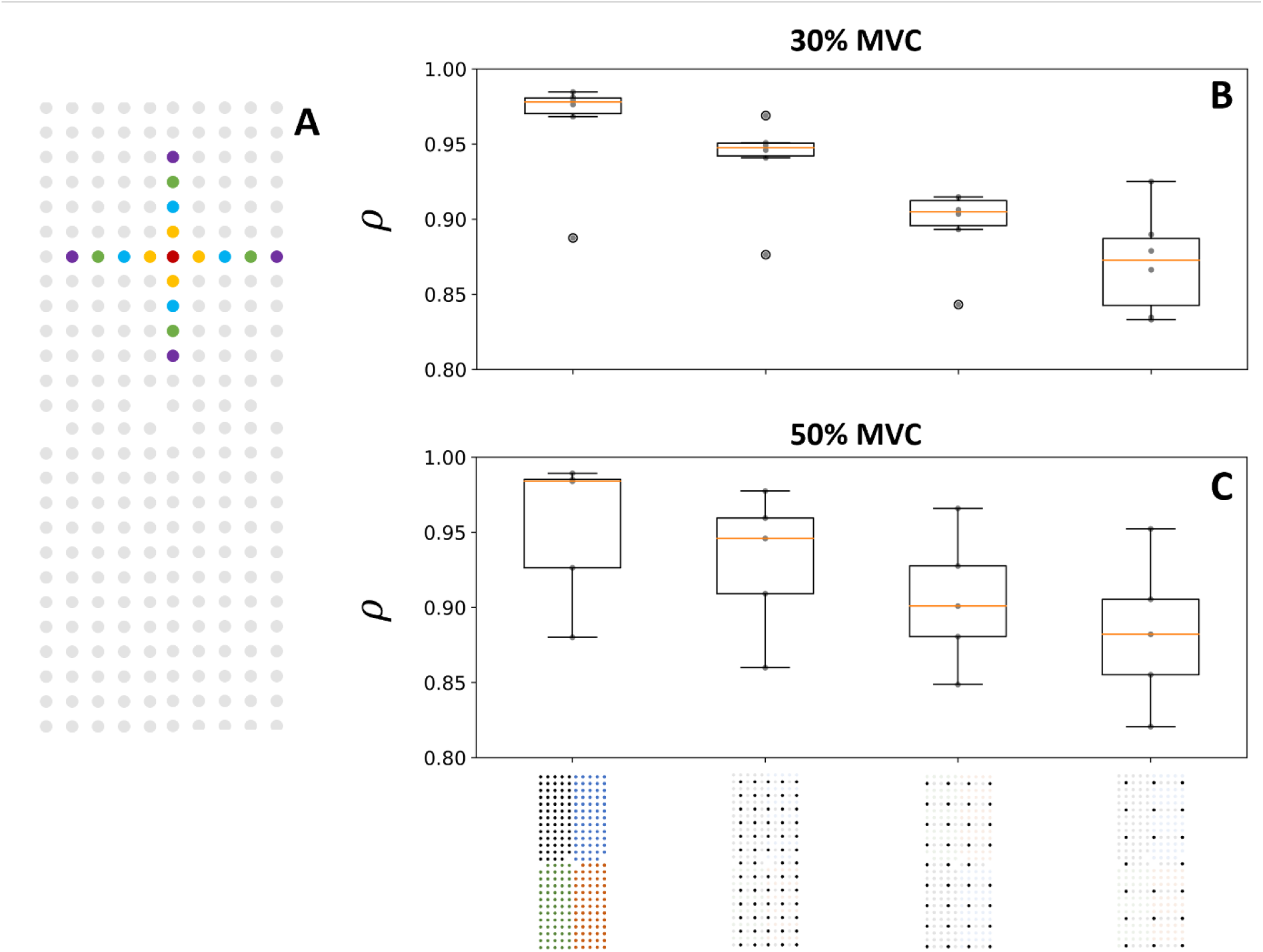
Effect of the electrode density on the correlation ρ between the profiles of motor unit action potentials (MUAP) detected over adjacent electrodes (A) at 30% (B) and 50% MVC (C). The profile of the MUAP detected over the red electrode was compared to those detected over the four adjacent electrodes separated by a 4 (orange), 8 (blue), 12 (green) and 16 (purple) mm IED (A). The boxplots denote the correlation coefficient ρ per participant (grey dots) and the median (orange line), quartiles, and 95%-range across participants.

### Laboratory study with an ultra-dense prototyped grid of 256 electrodes with 2-mm IED

31 and 26 motor units (PNR > 28 dB) were identified for one participant with the ultra-dense grid of 256 electrodes (2-mm IED, 9 cm^2^, Figure 1H) at 30% and 50% MVC, respectively (Figure 9B). Note that the signals from four independent subsets of 64 electrodes were decomposed separately. For that participant, more motor units were identified with the ultra-dense grid of 256 electrodes than with the grid of 64 electrodes covering the same area (Figure 5A, C). Indeed, 31 and 26 motor units were respectively identified at 30% and 50% MVC with the grid of 256 electrodes (Figure 9C), while 25 (24 ± 5 for the group) and 19 (18 ± 4 for the group) motor units were identified with the grid of 64 electrodes (Figure 5A, C). Moreover, fewer motor units were identified when the electrode density of the ultra-dense grid was decreased (Figure 9C), with 22 and 13 motor units identified with a 4- and 8-mm IED at 30% MVC, respectively, and 21 and 9 motor units identified with a 4- and 8-mm IED at 50% MVC, respectively. At 30% MVC, the rate of increase of N between 4- and 2-mm IED followed the prediction computed in Figure 4B and illustrated by the dash line in Figure 9C. At 50% MVC, the rate of increase of N (dotted line in Figure 9C) was lower than the prediction. As previously observed, the correlation between adjacent MUAPs increased from ρ = 0.92 with an 8-mm IED to ρ = 0.98 with a 2-mm IED at 30% MVC, and from ρ = 0.85 with an 8-mm IED to ρ = 0.93 with a 2-mm IED at 50% MVC (Figure 9A). All the motor units identified with the 8-mm and 4-mm IED were also identified with the 4-mm and 2-mm IED grids, respectively. Finally, more motor units with an early recruitment were identified when increasing the density from 8-to 4-mm IED (blue vs black trains in Figure 9B), and from 4-to 2-mm IED (red trains in Figure 9B).

**Figure 9:**
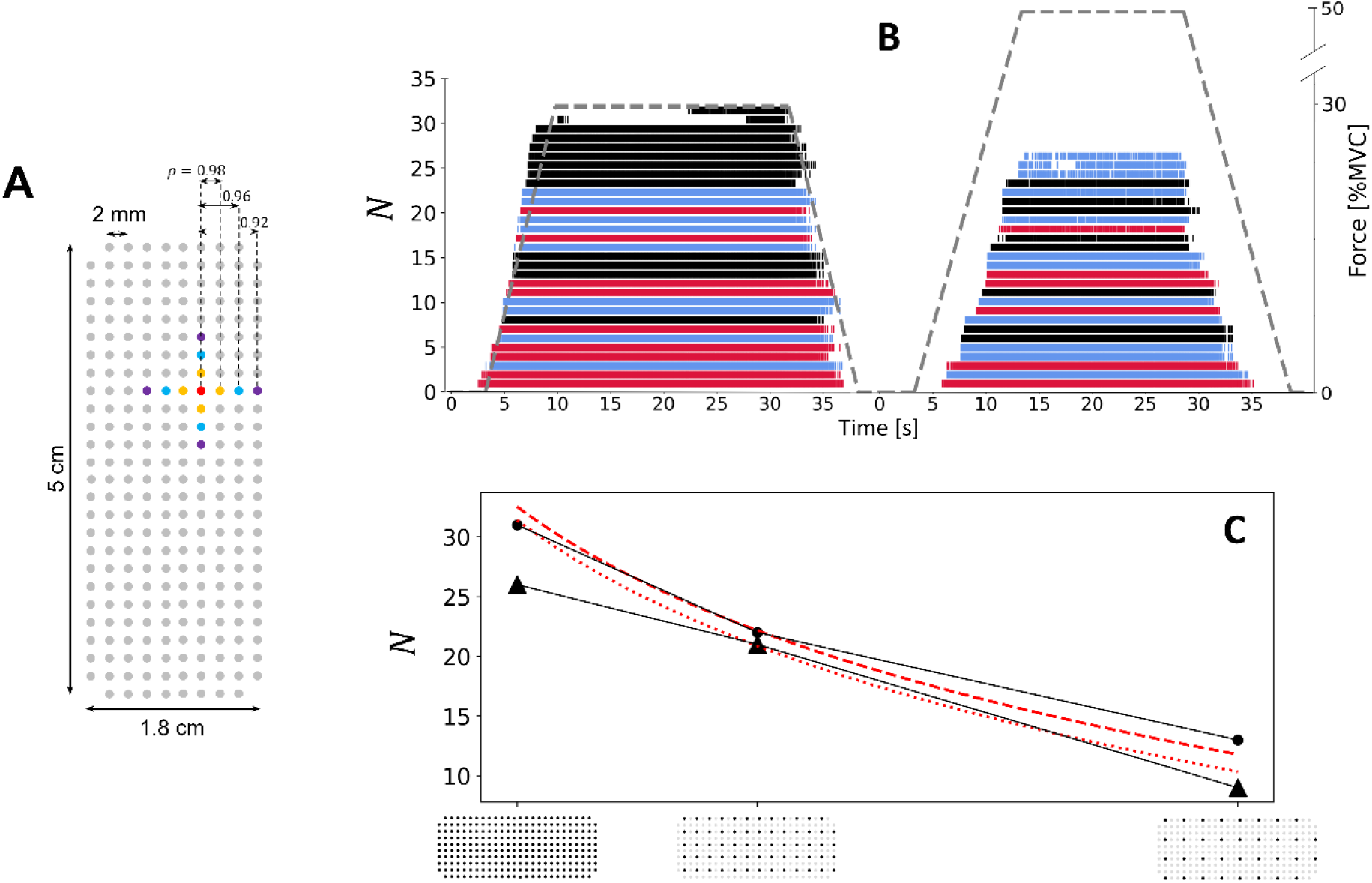
Results for the ultra-dense prototyped grid (2 mm IED, 5 x 1.8 cm, 256 electrodes). (A) Description of the ultra-dense grid, where grey circles represent the electrodes. On average, the correlation between the profiles of MUAPs detected over electrodes separated by an IED of 2 mm (orange), 4 mm (blue), and 8 mm (purple) reached ρ = 0.98, 0.96, and 0.92 at 30% MVC, respectively, and 0.93, 0.88, and 0.85 at 50% MVC, respectively. (B) Series of discharge times for motor units identified at 30% (left) and 50% MVC (right). The dark ticks represent the discharge times identified with a grid of electrodes with an 8-mm IED. The discharge times in blue were additionally identified with a grid of electrodes with a 4-mm IED, and the discharge times in red were additionally identified with a grid of electrodes with a 2-mm IED. All the pulse trains identified with one grid were also identified with the denser grids. (C) Effect of electrode density on the number of identified motor units at 30% (scatters) and 50% MVC (triangles). The trendlines from the density analysis inFigure 4B, D are also reported (red dotted lines). To maintain consistency with the other results, the grid was decomposed as four independent subsets of 64 electrodes, as explained in the Methods, to identify the higher number of motor units.

## Discussion

This study systematically investigated how the design parameters of grids of surface EMG electrodes (grid size and electrode density) impact the number and the properties of the motor units identified with EMG decomposition. Using a combination of computational and experimental analyses, we found that larger and denser grids of electrodes than conventionally used reveal a larger sample of identified motor units. As most of the motor units that were not identified with less dense and smaller grids had an early recruitment threshold, we concluded that denser grids allow to identify smaller motor units. This is due to a better spatial sampling of MUAPs over the grid, which in turn improves the discrimination of motor units with a unique set of MUAPs among active motor units. These results clarify the direction for designing new grids of electrodes that could span across the entire surface of the muscle of interest while keeping a high density of electrodes, with IED as low as 2 mm. Identifying large sets of small and large motor units is relevant in many research areas related to motor control, such as the investigation of synergies (Hug et al., 2022), neuromuscular modelling (Caillet et al., 2022c), or human-machine interfacing (Farina et al., 2021).

The number *N* of identified motor units increased across participants with the density of electrodes (Figure 4; Figure 8C), the size of the grid (Figure 6), and the number of electrodes (Figure 6). On average, 30 and 19 motor units were identified with the ‘conventional’ 64-electrode grid (8-mm IED, 32 cm^2^ surface area) at 30% and 50% MVC, respectively, which is consistent with several previous studies using similar grid designs (Del Vecchio et al., 2020). By increasing the density of electrodes and size of the grid to reach a total of 256 electrodes separated by a 4-mm IED, we identified on average 56 and 45 motor units at 30% and 50% MVC, respectively. We even reached 79 and 59 motor units for one subject (Figure 3), which is substantially more than the numbers of motor units usually reported in studies with similar methods, and twice those obtained with grids of 64 electrodes in this study. Our computational and experimental analyses showed that the size of the grid is a key factor contributing to the higher number of identified motor units (Figure 2B; Figure 6). According to our simulations, increasing the size of the grid increases the number of theoretically identifiable motor units, i.e., the number of motor units with unique sets of MUAPs across electrodes (Figure 2B). These differences between MUAPs result from the anatomical and physiological differences between adjacent motor units, such as the length of their fibers, the spread of the end plates, or their conduction velocity, as well as from the properties of the tissues separating the fibers from each recording electrode (Farina et al., 2004). Larger grids better sample these differences across electrodes, revealing the unique profiles of each motor unit action potentials (Farina et al., 2008). The density of electrodes was also a critical factor to increase the number of identified motor units (Figure 4; Figure 4-2C). Dense grids especially allowed to better identify early recruited motor units. Classically, the decomposition algorithms tend to converge towards the large and superficial motor units that contribute to most of the energy of the EMG signals (Farina and Holobar, 2016). Conversely, action potentials of the smallest motor units tend to have lower energy and are masked by the potentials of the larger units. These factors explain the lowest representation of low-threshold motor units in available HD-EMG datasets (Caillet et al., 2023). Increasing the density of electrodes would therefore enable to better sample the action potential profiles of these early recruited motor units across multiple electrodes, enabling their identification. However, we observed that increasing the density did not reveal additional early recruited motor units during contractions at 50% MVC (Figure 7D). This is potentially due to the higher energy of the MUAPs of the motor units recruited between 30% and 50% MVC. Additionally, we also showed in one subject that synthetically increasing the density of electrodes by resampling EMG signals with spatial interpolation does not have the same effect as with denser grids. In this example, 4 and 19 motor units were identified from the interpolated grid with a 4-mm and 2-mm IED, respectively, vs. 19 and 24 motor units with the experimentally recorded signals. All the motor units identified with the interpolated grid were also identified with the experimentally recorded signals (Figure 4-2).

The number of identified motor units *N* monotonically increased with the density of electrodes (Figure 4BD), the size of the grid (Figure 6BD) and the number of electrodes (Figure 6), following significant logarithmic trendlines. Remarkably, very similar logarithmic tendencies were obtained at both 30% and 50% MVC in all the analyses. Altogether, these trendlines suggested that the normalized number of identified motor units 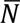 would grow with an electrode density beyond a 4-mm IED. We experimentally tested this hypothesis by designing a new prototyped grid of 256 electrodes separated by an IED of 2 mm. As predicted, more motor units were identified with a 2 mm than with a 4 mm IED, following at 30% MVC the same rate of increase as predicted by the logarithmic trendlines (Figure 4-2C) between 4-mm and 2-mm IED. This increase may plateau with higher electrode densities, as the level of correlation between the profiles of MUAPs detected over adjacent electrodes tended to 1 (Figure 4-2A). Therefore, the high level of similarity between signals recorded from adjacent electrodes in ultra-dense grids (IED < 2 mm) may limit the percentage of identifiable motor units (Farina and Holobar, 2016). According to these results, we consider that optimal designs of surface grids of electrodes for identifying individual motor units would involve a surface that covers the muscle of interest with an IED as low as 2 mm.

Another important factor for the accuracy of the discharge times estimated for each individual motor unit is the quality of the motor unit pulse trains, estimated by the PNR (Holobar et al., 2014) or the silhouette value. In this study, we found that the quality of the identified motor units (i.e., decomposition accuracy) increased when increasing the density of electrodes or the size of the grid, with PNR reaching on average 37-38 dB across participants with the grid of 256 electrodes (Figure 4-3). A greater average PNR implies the need of less manual editing following the automatic decomposition (Hug et al., 2021b). The better estimates of motor unit pulse trains depend on the better signal to noise ratio following the inversion of the mixing matrix, since the pulse train of each motor unit is computed by projecting the extended, whitened signals on the separation vector (Holobar and Farina, 2014; Farina and Holobar, 2016; Negro et al., 2016). Likewise, the PNR substantially increased after we computationally increased the number of electrodes by spatially resampling the EMG signals. This practical result is of interest for most of the physiological studies that require a lengthy processing time to visually inspect and manually edit the discharge times estimated from the pulse trains of all the motor units (Hug et al., 2021b).

Finally, we increased both the total number and the percentage of early recruited motor units identified by independently decomposing subsets of 64 electrodes within the grids of 256 electrodes, compared to the simultaneous decomposition of all available observations (Figure 7B, C). This was likely due to the lower ratio of large motor units sampled by each subset of electrodes, allowing the algorithm to converge to smaller motor units that contributed to the signal (Figure 7B, C). Importantly, it should be noted that the simulation results were obtained independently of a specific decomposition algorithm, as previously proposed by Farina et al (2008). On the other hand, the experimental results are based on a specific algorithm. Interestingly, however, the simulation and laboratory results were fully consistent and in agreement, indicating that the difference in shape of the spatially sampled MUAPs is the main factor influencing EMG decomposition.

## Conclusion

By increasing the density and the number of electrodes, and the size of the grids, we increased the number of theoretically identifiable and experimentally identified motor units from the surface EMG signals. The identified motor units had pulse trains with high PNR, limiting the manual processing time. Moreover, we identified a higher percentage of early recruited motor units, which are classically filtered out with the conventional grid designs. In this way, a maximum of 79 motor units (PNR > 28 dB; mean: 36 dB), including 40% of early recruited motor units, were identified, which is substantially greater than the samples previously reported with smaller and less dense grids. From these results, we encourage researchers to develop and apply larger and denser EMG grids to cover the muscle of interest with IEDs as small as 2 mm. This approach should increase the sample of motor units that can be experimentally investigated with non-invasive techniques.

## Notes

### Competing Interest Statement

The authors have declared no competing interest.

### Summary of Updates

Edited some phrasing Extended the Methods section on HDEMG decomposition

## References

Avrillon S, Hug F, Gibbs C, Farina D (2023) Tutorial on MUedit: An open-source software for identifying and analysing the discharge timing of motor units from electromyographic signals. bioRxiv:2023.2007.2013.548568.

Caillet AH, Phillips ATM, Farina D, Modenese L (2022a) Mathematical relationships between spinal motoneuron properties. Elife 11.

Caillet AH, Phillips ATM, Farina D, Modenese L (2022b) Estimation of the firing behaviour of a complete motoneuron pool by combining electromyography signal decomposition and realistic motoneuron modelling. PLoS Comput Biol 18:e1010556.

Caillet AH, Phillips ATM, Carty CP, Farina D, Modenese L (2022c) Hill-type computational models of muscle-tendon actuators: a systematic review. bioRxiv:2022.2010.2014.512218.

Caillet AH, Phillips ATM, Farina D, Modenese L (2023) Motoneuron-driven computational muscle modelling with motor unit resolution and subject-specific musculoskeletal anatomy. bioRxiv 2023.06.03.543552; doi: https://doi.org/10.1101/2023.06.03.543552

Churchland MM, Shenoy KV (2007) Temporal complexity and heterogeneity of single-neuron activity in premotor and motor cortex. J Neurophysiol 97:4235–4257.

Del Vecchio A, Negro F, Felici F, Farina D (2017) Associations between motor unit action potential parameters and surface EMG features. J Appl Physiol (1985) 123:835-843.

Del Vecchio A, Holobar A, Falla D, Felici F, Enoka RM, Farina D (2020) Tutorial: Analysis of motor unit discharge characteristics from high-density surface EMG signals. J Electromyogr Kinesiol 53:102426.

Duchateau J, Enoka RM (2011) Human motor unit recordings: origins and insight into the integrated motor system. Brain Res 1409:42–61.

Farina D, Holobar A (2016) Characterization of Human Motor Units From Surface EMG Decomposition. Proceedings of the Ieee 104:353–373.

Farina D, Merletti R, Enoka RM (2004) The extraction of neural strategies from the surface EMG. J Appl Physiol (1985) 96:1486-1495.

Farina D, Negro F, Gazzoni M, Enoka RM (2008) Detecting the unique representation of motor-unit action potentials in the surface electromyogram. J Neurophysiol 100:1223–1233.

Farina D, Negro F, Muceli S, Enoka RM (2016) Principles of Motor Unit Physiology Evolve With Advances in Technology. Physiology (Bethesda) 31:83–94.

Farina D, Vujaklija I, Brånemark R, Bull AMJ, Dietl H, Graimann B, Hargrove LJ, Hoffmann KP, Huang HH, Ingvarsson T, Janusson HB, Kristjánsson K, Kuiken T, Micera S, Stieglitz T, Sturma A, Tyler D, Weir RFF, Aszmann OC (2021) Toward higher-performance bionic limbs for wider clinical use. Nat Biomed Eng.

Gallego JA, Perich MG, Chowdhury RH, Solla SA, Miller LE (2020) Long-term stability of cortical population dynamics underlying consistent behavior. Nat Neurosci 23:260–270.

Heckman CJ, Enoka RM (2012) Motor unit. Compr Physiol 2:2629–2682.

Henneman E, Mendell LM (1981) Functional Organization of Motoneuron Pool and its Inputs. In: Comprehensive Physiology, pp 423–507.

Holobar A, Farina D (2014) Blind source identification from the multichannel surface electromyogram. Physiol Meas 35:R143–165.

Holobar A, Minetto MA, Farina D (2014) Accurate identification of motor unit discharge patterns from high-density surface EMG and validation with a novel signal-based performance metric. J Neural Eng 11:016008.

Holobar A, Minetto MA, Botter A, Negro F, Farina D (2010) Experimental analysis of accuracy in the identification of motor unit spike trains from high-density surface EMG. IEEE Trans Neural Syst Rehabil Eng 18:221–229.

Hug F, Del Vecchio A, Avrillon S, Farina D, Tucker K (2021a) Muscles from the same muscle group do not necessarily share common drive: evidence from the human triceps surae. J Appl Physiol (1985) 130:342-354.

Hug F, Avrillon S, Sarcher A, Del Vecchio A, Farina D (2022) Correlation networks of spinal motor neurons that innervate lower limb muscles during a multi-joint isometric task. J Physiol.

Hug F, Avrillon S, Del Vecchio A, Casolo A, Ibanez J, Nuccio S, Rossato J, Holobar A, Farina D (2021b) Analysis of motor unit spike trains estimated from high-density surface electromyography is highly reliable across operators. J Electromyogr Kinesiol 58:102548.

Jun JJ et al. (2017) Fully integrated silicon probes for high-density recording of neural activity. Nature 551:232–236.

Konstantin A, Yu T, Le Carpentier E, Aoustin Y, Farina D (2020) Simulation of Motor Unit Action Potential Recordings From Intramuscular Multichannel Scanning Electrodes. IEEE Trans Biomed Eng 67:2005–2014.

Muceli S, Poppendieck W, Holobar A, Gandevia S, Liebetanz D, Farina D (2022) Blind identification of the spinal cord output in humans with high-density electrode arrays implanted in muscles. Science advances 8:eabo5040.

Muceli S, Poppendieck W, Negro F, Yoshida K, Hoffmann KP, Butler JE, Gandevia SC, Farina D (2015) Accurate and representative decoding of the neural drive to muscles in humans with multi-channel intramuscular thin-film electrodes. J Physiol 593:3789–3804.

Negro F, Muceli S, Castronovo AM, Holobar A, Farina D (2016) Multi-channel intramuscular and surface EMG decomposition by convolutive blind source separation. J Neural Eng 13:026027.

Steinmetz NA, Koch C, Harris KD, Carandini M (2018) Challenges and opportunities for large-scale electrophysiology with Neuropixels probes. Curr Opin Neurobiol 50:92–100.

Stringer C, Pachitariu M, Steinmetz N, Reddy CB, Carandini M, Harris KD (2019) Spontaneous behaviors drive multidimensional, brainwide activity. Science 364:255.

